# Nasopharyngeal Microbiota as an early severity biomarker in COVID-19 hospitalised patients: a retrospective cohort study in a Mediterranean area

**DOI:** 10.1101/2021.09.28.461924

**Authors:** Maria Paz Ventero, Oscar Moreno-Perez, Carmen Molina-Pardines, Andreu Paytuví-Gallart, Vicente Boix, Irene Galan, Pilar González-delaAleja, Mario López-Pérez, Rosario Sánchez-Martínez, Esperanza Merino, Juan Carlos Rodríguez

## Abstract

**Background:** There is mounting evidence suggesting that the microbiome composition could be different in COVID-19 patients. However, the relationship between microbiota and COVID-19 severity progression is still being assessed. This study aimed to analyse the diversity and taxonomic composition of the nasopharyngeal microbiota, to determine its association with COVID-19 clinical outcome.

**Methods and Findings:** Samples came from a retrospective cohort of adult patients with COVID-19, hospitalised in a tertiary centre. To study the nasopharyngeal microbiota, we utilized 16S rRNA sequencing. Raw sequences were processed by QIIME2. The associations between the microbiota, invasive mechanical ventilation (IMV), and all-cause mortality were analysed by multiple logistic regression (OR; 95%CI), adjusted for age, gender, and comorbidity. 177 patients were included: median age 68.0 years, 57.6% males, 59.3% had a Charlson comorbidity index ≥3, and 89.2% with pneumonia. The microbiota α diversity indexes were lower in patients with a fatal outcome, and this association persisted after adjustment for the main confounders; whereas the β diversity analysis showed a significant clustering, grouping the patients with a fatal outcome. After multivariate adjustment, the presence of *Selenomonas spp*., *Filifactor spp*., *Actinobacillus spp*., or *Chroococcidiopsis spp*., was associated with a reduced risk of IMV (adjusted OR 0.06[95%CI 0.01–0.0.47], p = 0.007).

**Conclusions:** The microbiota diversity and taxonomic composition are related to COVID-19 severity. Higher diversity and the presence of certain genera in the nasopharyngeal microbiota seem to be early biomarkers of a favourable clinical evolution in hospitalised patients with moderate to severe SARS-CoV-2 infections.

## Introduction

In this time of pandemic finding early prognostic markers of COVID-19 severity is of utmost importance [1,2]. It is known that poor outcomes related to COVID-19 are not only a consequence of the viral infection, but are also related to an aberrant host immune response, including the vast release of cytokines by the immune system, leading to uncontrolled inflammation and multi-organ failure [3].

Several risk or prognostic factors, such as genetic factors, comorbidities, age, sex, and geographical location, have been associated with COVID-19 severity [2,4,5]. Taken together, these characteristics could have a determining role in promoting immune responses, and preventing an excessive anti-viral immune reaction.

Microbiota may be related to or influence the natural history of certain infectious diseases [6]. For example, in *Clostridioides difficile* infection, a lower diversity of microbiota and a decrease in several families are associated with the incidence and clinical evolution of the disease [7]. Likewise, the respiratory microbiota have also been correlated with the clinical evolution of chronic respiratory diseases [8] and respiratory viral infections [9].

Regarding microbiota and COVID-19 pathology, many published studies have focused on the differences between COVID-19 and non-COVID-19 patients, suggesting a possible role of the gut or respiratory microbiota in susceptibility to SARS-CoV-2 infection [10,11]. Additionally, some studies have shown a relationship between the composition of the gut and respiratory microbiota and disease severity [12]. This relationship appears to be mainly based on the capacity of the microbiota to modulate the immune response [13,14], through modification of the gut-lung axis [12,15,16], and to alter the expression of angiotensin-converting enzyme 2 (ACE2) receptors, which are used by SARS-CoV-2 to enter host cells [17,18].

The available evidence suggests a potential role of microbiota in susceptibility to SARS-CoV-2 infection and COVID-19 severity, but longitudinal studies evaluating the microbiota as a prognostic factor for severity of disease progression are lacking. The data regarding the association between nasopharyngeal microbiota features and disease severity are scarce, and limited in terms of showing a decrease in α diversity or identifying specific genera with relevance to critical illness [19,20]. Since the sampling of this location is very accessible, with the nasopharyngeal aspirate swab diagnostic confirmation procedure able to obtain this information, it should be a priority to address the relationship between nasopharyngeal microbiota and COVID-19 outcomes.

This study aimed to analyse the nasopharyngeal microbiota from hospitalised COVID-19 patients, to determine the relationship between the microbiota and SARS-CoV-2 infection clinical outcomes and to identify features or genera that could be used as severity prognostic markers.

## Materials and methods

### Patients and study design

A retrospective cohort of adult patients with COVID-19, hospitalised in a tertiary centre (Alicante University General Hospital, Spain) from February 27^th^ 2020 to January 22^nd^ 2021, was studied. SARS-CoV-2 infection was confirmed by the RT-PCR-COBAS 6800 System (Roche Molecular Systems, Branchburg, NJ, United States). At hospital admission one nasopharyngeal specimen per patient was obtained, stored at -80°C and later analysed.

Of the 1526 patients hospitalised in the study period, nasopharyngeal samples from 324 patients were randomly processed and preserved. Due to the available economic resources, sixty percent of the samples were randomly sampled for processing; 17 samples did not correspond to the first PCR sample, so they were discarded. Finally, 177 patients were included in the study.

### Variables and data collection

The clinical features, comorbidity, laboratory and radiological tests, prescribed therapies, and outcome during the acute phase of the infection by SARS-CoV-2 were extracted from the digital medical record.

The main explanatory variables of the analysis were the microbiota diversity, measured by the α and β diversity indexes, and the taxonomic composition, expressed by the differentially represented genera.

#### Primary Outcomes

Invasive mechanical ventilation (IMV) and all-cause mortality.

### DNA isolation and microbiota amplicon next-generation sequencing (NGS)

The nasopharyngeal samples frozen at –80 °C were used for DNA isolation with the QIAamp MiniElute Virus Spin Kit (Quiagen, Hilden, Germany), following the protocol recommended by the manufacturer. The DNA obtained was quantified with a Qubit 4 Fluorometer, using a Qubit dsDNA HS Assay Kit (ThermoFisher Scientific, Massachusetts, United States). The microbiota amplicon sequencing was performed following the protocol of the 16S Metagenomics Sequencing Library Preparation recommended by Illumina. The V3 and V4 region from 16S rRNA gene were amplified by PCR, and then the fragments obtained were sequenced in the MiSeq system with V3 reagents (600 cycle, 2×300bp).

### Bioinformatic analyses

The raw reads obtained from the NGS were analysed using QIIIME2 (2021.2 version) [21]. The denoising was performed with the plugin DADA2 and to avoid contamination and false positives a BLAST against the database of human genome of NCBI was performed, as well as singletons were removed. The taxonomy was assigned using the SILVA Database (Release 132) [22]. Regarding the microbiota analyses, the Shannon, Pielou, and Simpson indexes were calculated to study the α diversity, and the UniFrac weighted distance plus PCoA were performed to analyse the β diversity. The genera that were differentially represented between severity groups (main outcomes present or not) were determined using the R package DESeq2 (4.1.0 version) [23]. The linear model obtained by DESeq2 was adjusted by the prescription of antibiotic treatment 3 months earlier.

### Statistical analysis

Categorical and continuous variables are given as frequencies (percentages) and as the median (interquartile range), respectively. Patients of the global cohort that were included and excluded were compared by Mann-Whitney’s U, chi-squared, and Fisher’s exact tests. Cumulative incidences of outcomes (95% confidence intervals (95%CI)) were registered. The final date of follow-up was March 1, 2021, unless censored. The differences between groups in the β diversity were assessed using the PERMANOVA test. Associations were evaluated by a chi-squared test. Multiple logistic regression models adjusted for age, gender, and comorbidity were built to evaluate the association between microbiota diversity indexes or the differentially represented genus (obtained by DESeq2) with the primary outcomes, and the odds ratios (OR) with the 95%CI were estimated. IBM SPSS Statistics v25 (Armonk, NY) was used for the analyses. P <0.050 defined statistical significance.

### Ethics statement and data availability

This project was performed in the Clinical and Biomedical Research Institute of Alicante (ISABIAL), under the written approval of the Ethics Committee of Clinical Research with Drugs (in Spanish, CEIm) of the General University Hospital of Alicante (Ref CEIm approval: PI2020-052).

The raw data from the sequencing are available in the National Center for Biotechnology Information Database (NCBI), under the Bioproject accession number PRJNA754005.

## Results

A total of 177 patients were included in the study. The study population and the global cohort of 1526 patients hospitalised while the study lasted were similar in age, gender, comorbidities, extent of infiltrates on chest radiograph, dexamethasone use, duration of hospitalization, and outcomes: IMV and mortality (p > 0.05).

Table 1 shows the general characteristics of the study population and the main features of the COVID-19 acute phase infection and its clinical evolution. The patients had a median age of 68.0 years (IQR) (52.0–80.0); 57.6% were males and 59.3 % had a Charlson comorbidity index ≥3. They were assessed in the emergency department after a median of 6 [3–7] days of symptoms, and 89.2% had pneumonia. Fifty-one patients (28.8%) had received antibiotic therapy in the 3 months prior to their hospital admission, for a median of 5 [2–6] days. The mortality rate was 17.5% (95%CI, 12.6–23.7) (31/177), and 11.3% (95%CI, 7.4–16.8) (20/177) required IMV.

**Table 1.** Demographic characteristics, comorbidities, clinical presentation, and clinical outcomes.

|  | Population<br>[n= 177] |
| --- | --- |
| **Demographics** |  |
| Age, median (IQR), years | 68 (52–80) |
| Age≥ 65 years old, % (N) | 55.9 (99/177) |
| Males, % (N) | 57.6 (102/177) |
| Nosocomial, % (N) | 1.7 (3/177) |
| Long-term care resident, % (N) | 4 (7/177) |
| Health professional, % (N) | 4 (7/177) |
| Waves<br>First (1.02.2020 - 31.05.2020), % (N)<br>Second (1.06.2020 - 15.12.2020), % (N)<br>Third (16.12.2020 - 31.03.2021), % (N) | 54.2 (96/177)<br>31.1 (55/177)<br>14.7 (26/177) |
| Antibiotic therapy in the previous 3 months | 28.8 (51/177) |
| **Comorbidities** |  |
| Hypertension, % (N) | 55.9 (99/177) |
| Diabetes, % (N) | 26.6 (47/177) |
| Current or former Smoker, % (N) | 20.6 (70/177) |
| Obesity, % (N) | 39.7 (56/141) |
| Chronic respiratory disease, % (N) | 21.6 (38/177) |
| Immunosuppression, % (N) | 4 (7/177) |
| Charlson comorbidity index, median (IQR) | 3 (1–6) |
| Charlson index $\geq 3$, % (N) | 59.3% (105/177) |
| 10-years expected survival<sup>a</sup> | 53.3 (1.6–90.1) |
| **Clinical Presentation** |  |
| Median time (IQR) from symptom to hospitalization, days<sup>b</sup> | 6 (3–7) |
| Fever, % (N) | 67.2 (119/177) |
| Cough, % (N) | 26.0 (46/177) |
| Dyspnoea, % (N) | 57.6 (102/177) |
| **Diarrhoea**, % (N) | 25 (44/177) |
| Confusion, % (N) | 9.6 (17/177) |
| Fatigue, % (N) | 41.0 (71/173) |
| Myalgias-arthralgias, % (N) | 30.1 (52/173) |
| Anosmia-dysgeusia, % (N) | 6.9 (12/173) |
| **Initial Assessment** |  |
| Oximetry $<94\%$ at room air, % (N) | 43.7% (73/167) |
| PaO<sub>2</sub>:FiO<sub>2</sub>, median (IQR) | 332 (272–404) |
| Respiratory rate, breaths/min, median (IQR) | 18 (16–24) |
| Systolic BP, mmHg, median (IQR) | 130 (118–145) |
| Diastolic BP, mmHg, median (IQR) | 78 (68–89) |
| Temperature, °C, median (IQR) | 36.9 (36.3–37.7) |
| Heart rate, beats/min, median (IQR) | 92 (81–102) |
| eGFR, ml/min/m<sup>2</sup>, median (IQR) | 73 (47–90) |
| Lymphocytes, per mm<sup>3</sup>, median (IQR) | 910 (700–1370) |
| Lymphopenia, % (N) | 44.3 (78/176) |
| C-reactive protein $> 10$ mg/dL, % (N) | 33.1 (55/175) |
| Procalcitonin $> 0.5$ ng/mL, % (N) | 12.4 (20/161) |
| Ferritin > 500 mg/L, % (N) | 59.8 (98/164) |
| Lactate dehydrogenase > 250 U/L, % (N) | 33.9 (53/156) |
| D-dimers > 1 mg/mL, % (N) | 33.1 (53/160) |
| Interleukin 6 ≥ 10 pg/mL, % (N) | 77.7 (101/130) |
| Troponin T > 14 ng/L, % (N) | 49.4 (77/176) |
| Brain natriuretic peptide > 125 pg/mL, % (N) | 53.5 (84/157) |
| Potassium mmol/L, median (IQR) | 4.1 (3.8–4.4) |
| Pneumonia on X-rays, % (N) | 89.2 (157/176) |
| Opacities >50% of lung surface on X-rays, % (N) | 21.5 (38/177) |
| **Treatment** |  |
| Corticosteroids, % (N) | 46.3% (82/177) |
| Remdesivir, % (N) | 3.9% (7/177) |
| Tocilizumab, % (N) | 23.7% (42/177) |
| **Outcomes** |  |
| Non-invasive mechanical ventilation requirement, % (N) | 23.1 (41/177) |
| Invasive mechanical ventilation requirement, % (N) | 11.3 (20/177) |
| Mortality, % (N) | 17.5 (31/177) |

Data shown as %, median (interquartile range, IQR), unless specified otherwise. In bold, statistically significant differences.
Percentages may not total 100 because of rounding.
<sup>a</sup>10-years expected survival derived from Charlson comorbidity index score.
<sup>b</sup>Days of symptoms before admission. OR: odds ratio, 95%CI: 95% confidence interval.

### Diversity analysis and outcomes

The α diversity indexes were lower in patients with a fatal outcome: Shannon 3.59[2.86– 4.42] vs. 4.39[3.12–5.14], p=0.014; Pielou 0.58[0.50–0.67] vs. 0.71[0.55–0.79], p=0.007; and Simpson index 0.80[0.62–0.88] vs. 0.89[0.76–0.94], p=0.018 (**Figs 1A, 1B**, and **1C**). The protective effect of a greater microbiota diversity persisted for the Shannon (adjusted OR (aOR) 0.654 [95%CI 0.448–0.956], p = 0.028) and Pielou indexes (aOR 0.055[95%CI 0.003–0.823], p = 0.036) after adjustment for age, gender, and comorbidities. The β diversity analysis showed a significant clustering (p= 0.014), grouping together the fatal outcome patients (**Fig 1D**). In the case of IMV, neither the α diversity indexes nor β diversity analyses showed any significant differences.

**Fig 1.**
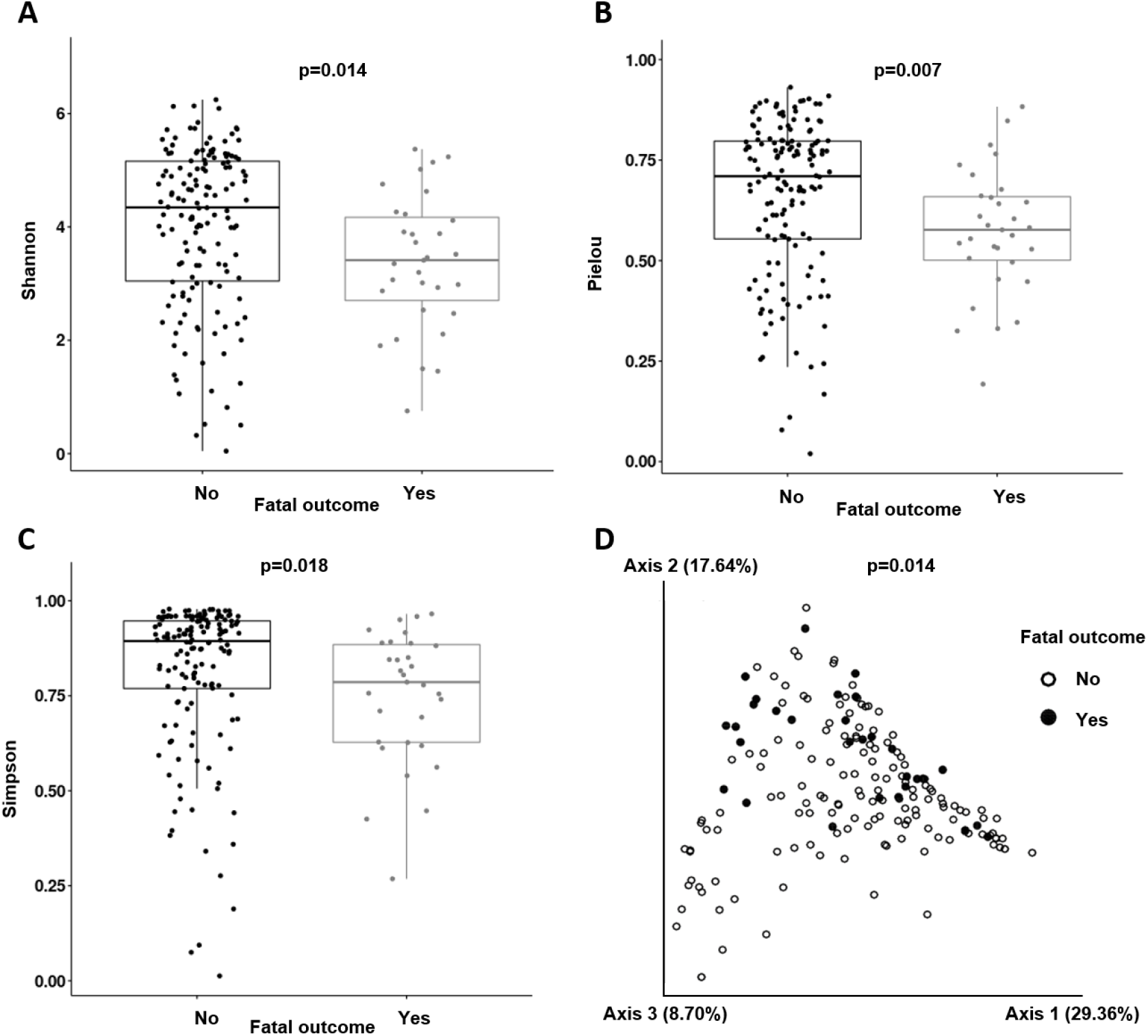
Diversity analysis: Boxplots obtained for the Shannon index (**A**), Pielou index (**B**), and Simpson index (**C**). PCoA (principal coordinates analysis) for the β diversity distribution along the samples (**D**).

### Taxonomic analysis and outcomes

*Streptococcus spp*. (14.14 %), *Staphylococcus spp. (12*.*12%)*, and *Corynebacterium spp. (9*.*11%)* were the genera that were more abundant in COVID-19 patients, without significant differences between patients with IMV or a fatal outcome. By group, there were 34.20% (483/1412) taxa shared between IMV/non-IMV subpopulations, 4.67% (66/1412) taxa exclusively found in IMV patients, and 61.12% (863/1412) taxa only detected in non-IMV patients (**Fig 2A**).

**Fig 2.**
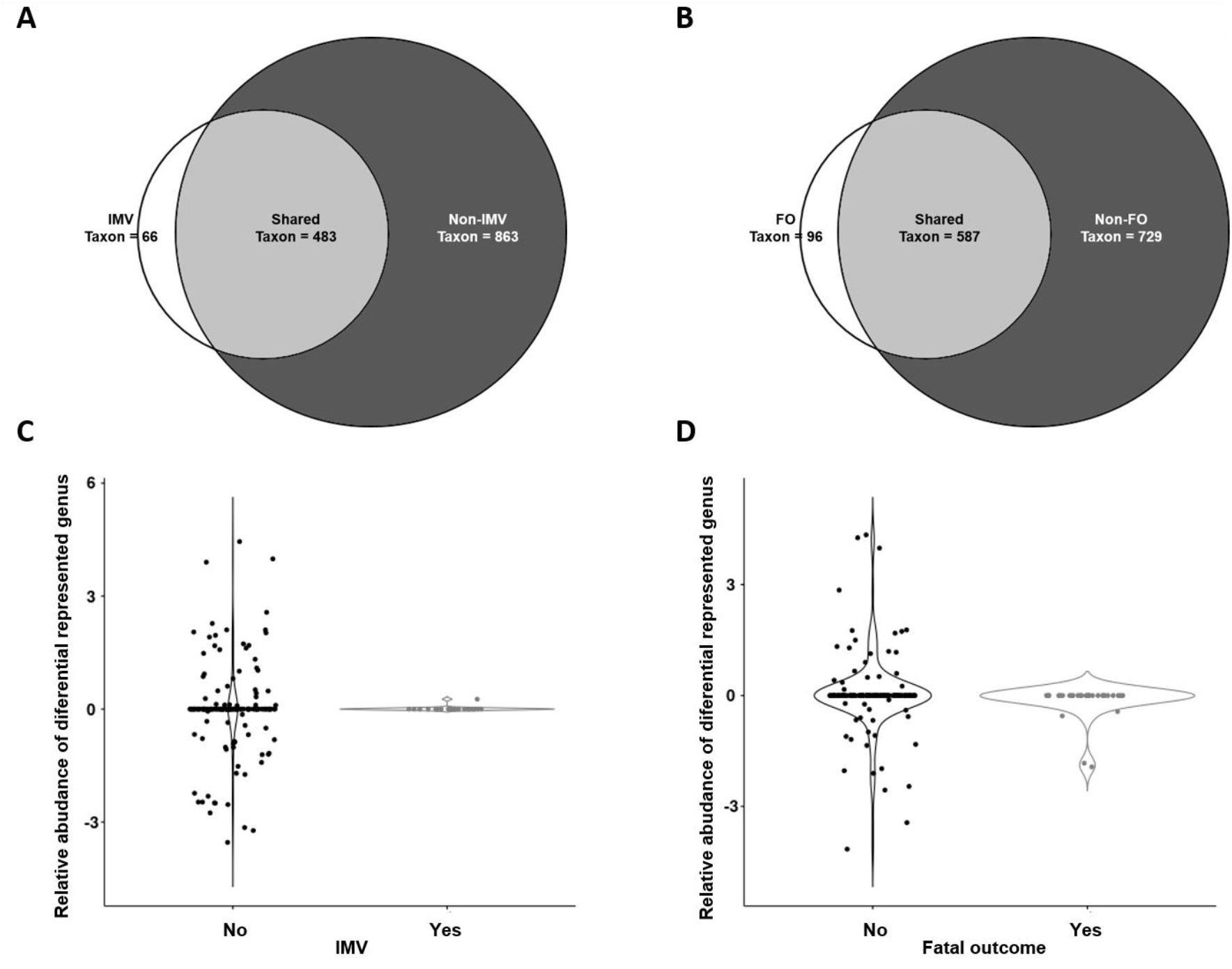
Taxonomic analysis: Venn diagrams for IMV (**A**), and fatal outcome (**B**), and relative abundances of differential genera for IMV (**C**), and fatal outcome subpopulations (**D**). Relative abundances are shown in logarithmic scale. IMV: invasive mechanical ventilation, FO: Fatal outcome.

Regarding fatal outcomes, the results were similar. The shared taxa comprised 41.57% (587/1412), taxa exclusively found in the exitus subpopulation were 6.8% (96/1412), and in survivors 51.2% (729/1412) (**Fig 2B**).

### Differently represented genera and outcomes

This study was performed to identify differential genera between the subpopulations with and without specific outcomes. We found that *Selenomonas spp. (LogFC= 23*.*96; p<0*.*0001), Filifactor spp. (LogFC= 23*.*51; p<0*.*0001), Actinobacillus* spp. *(LogFC= 24*.*86; p<0*.*0001)* and *Chroococcidiopsis spp. (LogFC= 22*.*31; p<0*.*0001)* were significantly more abundant in non-IMV patients (**Fig 2C**). The presence of *Selenomonas spp*., *Filifactor spp*., *Actinobacillus spp*., or *Chroococcidiopsis spp*., was associated with a reduced risk of IMV (OR 0.062 [95%CI 0.01–0.47], p = 0.007). This protective association persisted after adjustment for the main confounders in the multivariate model (**Fig 3**).

**Fig 3.**
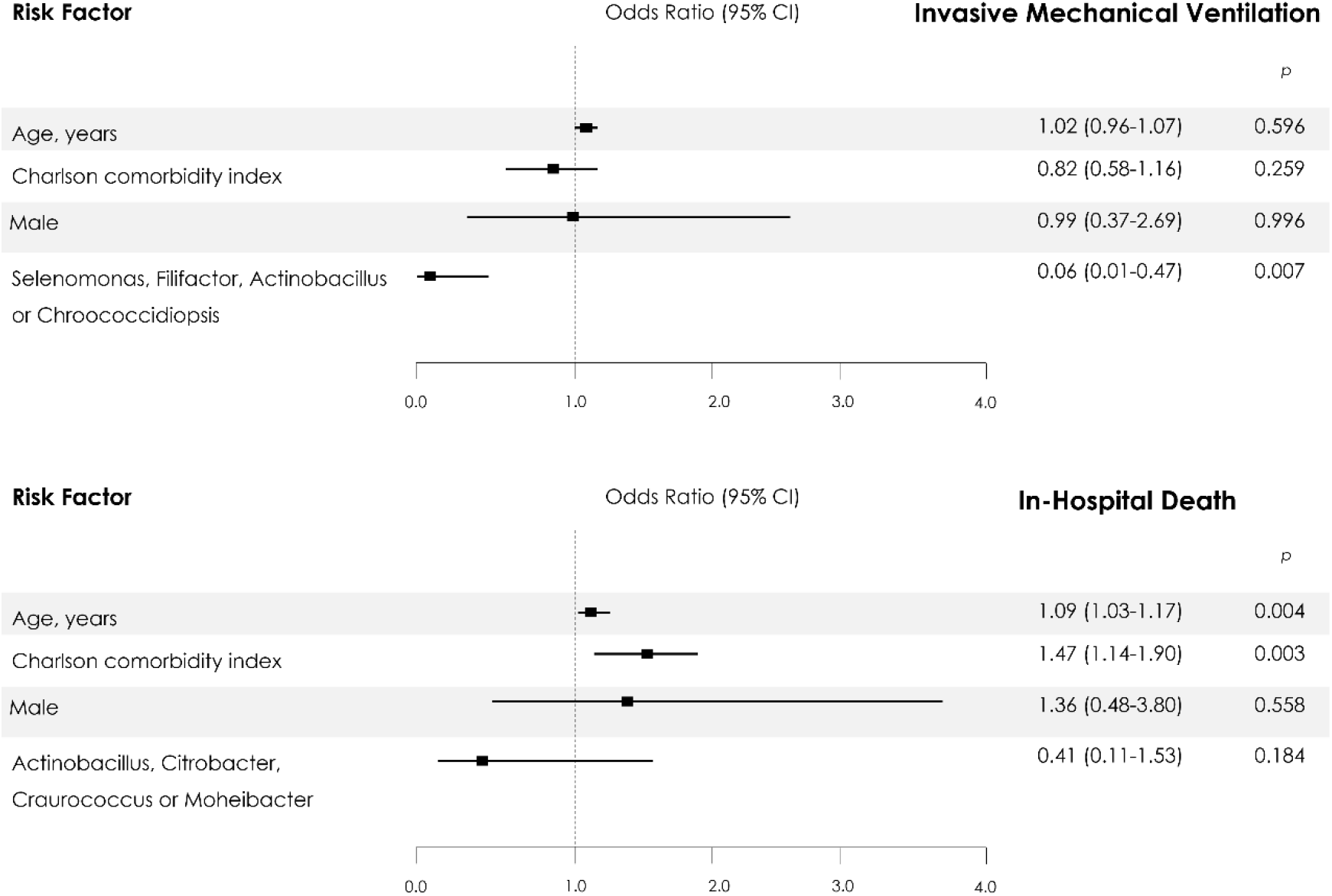
Predictors of Invasive Mechanical Ventilation and In-Hospital Death from Multivariable Logistic-Regression Analysis. The 95% confidence intervals (CIs) of the odds ratios have been adjusted for multiple testing.

For fatal outcomes, *Actinobacillus spp. (LogFC= 24*.*30; p<0*.*0001), Citrobacter spp. (LogFC= 25*.*21; p<0*.*0001), Craurococcus spp. (LogFC= 22*.*77; p<0*.*0001)*, and *Moheibacter spp. (LogFC= 22*.*7; p<0*.*0001)* were significantly more abundant in non-exitus patients (**Fig 2D**). The presence of *Actinobacillus spp*., *Citrobacter spp*., *Craurococcus spp*., *or Moheibacter spp*., was associated with a reduced risk of a fatal outcome (OR 0.309[95%CI 0.10–0.93], p = 0.037). This association did not persist after adjustment for the main confounders in the multivariate model (**Fig 3**).

## Discussion

Recently, several studies assessing the relationship between the gut microbiome and the severity of COVID-19 have been published [24,25]. However, to our knowledge, this is the first study that has evaluated nasopharyngeal microbiota at the time of admission as a prognosis biomarker of severity of disease progression in the acute infection phase of SARS-CoV-2, in a large cohort of hospitalised patients with COVID-19. The assessment showed a significant decrease of all diversity indexes studied (Shannon, Pielou, and Simpson) in patients with a final fatal outcome, linking an initial low microbiota diversity with COVID19 severity. The presence of four specific genera, *Selenomonas spp*., *Filifactor spp*., *Actinobacillus spp*. or *Chroococcidiopsis* spp., was associated with a reduction of more than 90% of IMV, regardless of age, gender, or comorbidity. The presence of *Actinobacillus spp*., *Citrobacter spp*., *Craurococcus spp*. or *Moheibacter spp*. was associated with a 70% reduction in mortality, but this relationship did not persist after adjustment for the main confounders.

The relationship between the microbiota and COVID-19 is an active and expanding field of research. Previous studies have been focused in the differences of the gut microbiota between COVID-19 and non-COVID19 patients, or its correlation with severity inflammatory markers [10,11]. However, there has been limited investigation into the relationship between microbial communities and COVID-19 clinical outcome.

Regarding COVID-19 and the gut microbiome, Gu et al. [26] reported that COVID-19 patients had a lower diversity microbiota (Shannon and Chao1 index) than healthy controls; also, several microorganisms (*Streptococcus spp*., *Rothia spp*., *Veillonella spp*. and *Actinomyces spp*.) were identified that could be used as COVID-19 biomarkers. According to these data, Zuo et al. [27], using the Bray-Curtis dissimilarities test, described alterations in the gut microbiome at the whole genome level, since their COVID19 patients were more heterogeneous than healthy controls. Yeoh et al. [12] found that specific genera, such as *Bifidobacterium adolescentis, Eubacterium rectale*, and *Faecalibacterium prausnitzii*, were depleted in the COVID-19 cohort when compared with non-COVID-19 patients, and were negatively correlated with the inflammatory marker CXCL10. The same correlation was reported by Zou et al. [27]. Likewise, Gou et al. [28] showed that the *Bacteroides* genus, and specifically *B. ovatus*, was associated with inflammatory cytokines such as IL-6, TNF-α and IFN-γ [28]. These depleted species in COVID-19 patients are known to play immunomodulatory roles in the human gastrointestinal system [29].

In terms of the association of the upper respiratory tract microbiome and SARS-COV-2 infection, the studies performed to date have included small cohorts of patients. Braun et al. [30] (n=33), De Maio et al. [31] (n=40), and Liu et al. [32] (n=9) showed no significant differences in the nasopharyngeal microbial community between COVID-19 and control patients using α-β diversity and taxonomic compositional analysis. Whereas Mostafa et al. [33] (n=50) and Engen et al. [34] (n=19) reported a lower α diversity (Chao1, Shannon, and Simpson indexes) in COVID-19 compared to healthy patients, and both groups showed significant dissimilarities in β diversity. Therefore, there is controversy regarding lung and nasopharyngeal microbiota composition on SARS-CoV2 infection.

Regarding microbiota and COVID-19 severity, Ma et al. [19] explored the oropharyngeal microbiome in COVID-19 patients (n=31) with various severities (mild, moderate, severe, or critical) compared with flu patients (n=29) and healthy controls (n= 28) using high-throughput metagenomics. They showed that critical COVID-19 patients presented with a significant diminution in α diversity (Shannon index), while noncritical patients exhibited no significant change from the normal group.

The present work pioneered the analysis of the nasopharyngeal microbiota (using 16S rRNA gene sequencing), in a large cohort of hospitalised patients with COVID-19, as a prognosis biomarker. The lower diversity in patients with a fatal outcome is in agreement with the hypothesis that low microbiota diversity is associated with the development of several pathologies [35,36], and high diversity is associated with lower severity [37].

A study performed with 24 critically ill COVID-19 patients and 24 non-COVID-19 patients with pneumonia [38] showed taxonomical differences between the lung microbiota of COVID-19 and non-COVID-19 patients. The characteristic microorganisms of COVID-19 patients were *Pseudomonas alcaligenes, Sphingobacterium spp*., *Clostridium hiranonis* and *Acinetobacter schindleri*. While the genera that characterised the lung microbiota in the COVID-19-negative patients were *Streptococcus* spp., *Haemophilus* or *Selenomonas* spp. Regarding the upper respiratory tract microbiota, Ma et al. [19] found increased ratios of *Klebsiella* sp., *Acinetobacter* sp., and *Serratia* sp. were correlated with both disease severity and elevated systemic inflammation markers (neutrophil– lymphocyte ratio). Along the same lines, *Prevotella* spp. was also linked to COVID-19 severity, which has been hypothesised to suggest a possible relationship with the inflammatory response [20].

Our taxonomic analysis identified several microorganisms, such as *Selenomonas, Filifactor, Actinobacillus*, and *Chroococcidiopsis SAG 2023*, related to IMV, and *Craurococcus, Actinobacillus, Citrobacter* and *Moheibacter* related to a fatal outcome. Future research to determine their roles in COVID-19 development and evolution is required.

Our study has several limitations, this was an observational, retrospective, single-centre study, and collection of data was not standardized in advance. The sample size and the absence of differences in the characteristics of the global cohort of patients admitted to our hospital during the duration of the study reinforce the present data. Multiple factors can condition changes in microbiota, including the use of antibiotics. Nonetheless, the design of the statistical analysis adjusted for the use of antibiotic therapy in the 3 months prior to the inclusion of the study, allowing us to limit this bias. The exclusion of these patients from the study would have greatly limited the external validity of our results. Finally, the 16S ribosomal RNA amplicon sequencing approach to study the microbiota could introduce bias in the obtained data because this method does not allow the study of the whole microbiome, but only the genera amplified by PCR. Nevertheless, it is the most common technique to study microbiota in clinical samples. Moreover, the microbiota bioinformatics analysis has not been standardized yet, which hampered comparison interpretations of our results.

In summary, the higher diversity found in patients without IMV or a fatal outcome, together with the presence of certain genera in the nasopharyngeal microbiota, seemed to be an early biomarker of a favourable clinical evolution in a cohort of Mediterranean hospitalised patients with SARS-CoV-2 infection. Our findings have potential clinical relevance due to the feasibility and low cost of developing rapid molecular techniques to evaluate the diversity and detect these genera at the time of admission. These data, taken together with other prognostic markers already being implemented, may allow identifying patients with a good prognosis (i.e., a 70–90% reduction in unfavourable clinical outcomes). Considering the clinical significance of these findings and the ease of their application in daily practice, further investigation to confirm these data could be very relevant for improving COVID-19 management.

